# B1a cells protect against *Schistosoma japonicum*–induced liver inflammation and fibrosis by controlling monocyte infiltration

**DOI:** 10.1101/420083

**Authors:** Liang Yong, Yuanyuan Tang, Cuiping Ren, Miao Liu, Jijia Shen, Xin Hou

## Abstract

During *Schistosoma* infection, lack of B cells results in more severe granulomas, inflammation, and fibrosis in the liver, but the mechanisms underlying this pathology remain unclear. Thus, our aim was to clarify the mechanisms underpinning the immunomodulation of B cells in mice infected with *Schistosoma japonicum*. We found that B cell deficiency led to aggravated liver pathology, as demonstrated by increases in the size of the egg-associated granulomas, alanine transaminase levels, and collagen deposition. Compared with infected wild-type mice, infected B cell–deficient μMT mice showed increased infiltration of Ly6C^hi^ monocytes and higher levels of proinflammatory cytokines (tumor necrosis factor alpha, interleukin 6, and interleukin 12) and chemokines ([C-C motif] ligands (CCL)2, CCL3, CCL4, and CCL5). The results of flow cytometric analysis and cell transfer experiments showed that B1a cells increased significantly in the liver following *S. japonicum* infection, with some of those cells deriving from the peritoneal cavity. We also found that secretion of IL-10 from hepatic B cells increased significantly in infected wild-type mice and that this IL-10 was mainly derived from B1a cells. In addition, adoptively transferring peritoneal cavity B cells purified from wild-type, but not from IL-10–deficient mice, to μMT mice significantly reduced liver pathology and liver infiltration of Ly6C^hi^ monocytes. These reductions were accompanied by decreases in the expression levels of chemokines and inflammatory cytokines. Taken together, these data indicated that after *S. japonicum* infection, an increased number of hepatic B1a cells secrete IL-10, which inhibits the expression of chemokines and cytokines and suppresses the infiltration of Ly6C^hi^ monocytes into the liver thereby alleviating liver early inflammation and late fibrosis. Understanding this immunomodulatory role of B1a cells in schistosomiasis may lead to the development of therapeutic strategies for *Schistosoma*-induced liver disease.

**Author summary:** Infection with *Schistosoma,* a waterborne parasitic flatworm (trematode) commonly called a blood fluke, results in strong granulomatous inflammation caused by the deposition of eggs in the liver. A granuloma is a substantial immune cell infiltration around the eggs intermixed with liver cells that can protect the host against liver damage. However, excessive infiltration and inflammation can lead to severe liver injury and fibrosis. Here, we found that B1a cells accumulate in the liver of mice after *S. japonicum*–induced infection and that these B1a cells release the anti-inflammatory cytokine interleukin 10 to regulate inflammation. The B1a cell–derived interleukin 10 inhibits the expression of chemokines (which attract cells such as monocytes to sites of infection or inflammation) and thus restrains excessive infiltration of Ly6C^hi^ monocytes (which may have proinflammatory activity) into the liver, thereby alleviating early inflammation and later fibrosis. Our study provides insight into the immunomodulation of B1a cells in schistosomiasis and offers key information for the development of therapeutic strategies in *Schistosoma*-induced liver disease.

## Introduction

Schistosomiasis is a chronic disease with the characteristic pathological manifestation of granulomatous lesions around parasitic eggs deposited in the liver and intestine. Granulomas are driven by a type 2 immune response and are a critical component in limiting the amount of tissue damage and preventing acute mortality [1]. However, granulomas may ultimately lead to liver fibrosis and sometimes to death in chronically infected hosts [2].

Macrophages are a major cellular component of granulomas. Both monocyte-derived and resident macrophages engage surrounding parasite eggs during infection [3, 4]. The recruitment of Ly6C^hi^ monocytes is the dominant mechanism for expanding macrophage populations in the *Schistosoma*-infected liver [3]. Recruitment of Ly6C^hi^ monocytes to inflammatory sites depends on the interactions between chemokine (C-C motif) ligands (CCLs) and their receptors (CCRs), including CCL2-CCR2, CCL1-CCR8, CCL3/4/5-CCR1/5, and CXCL10-CXCR3 interactions [5-8]. The number of Ly6C^hi^ monocytes in the livers of *Schistosoma*-infected CCR2-deficient (*Ccr2*^−/−^) mice is significantly reduced compared with that in infected wild-type (WT) mice [3]. Liver recruitment of Ly6C^hi^ monocytes has been documented in viral infection, sterile heat injury to the liver, and ischemia–reperfusion damage when Ly6C^hi^ monocyte–derived macrophages have an M1 or proinflammatory phenotype and aggravate liver injury and fibrosis by releasing proinflammatory and profibrotic factors [9, 10]. In schistosomiasis, Ly6C^hi^ monocytes in granulomas respond to T helper 2 (Th2)-cell derived interleukin (IL)-4 and IL-13 to exhibit an arginase 1–positive, resistin-like molecule alpha–positive and chitinase-like 3–positive M2 phenotype or an alternatively activated macrophage (AAM) phenotype [11]. Studies using animal models have indicated that AAMs are essential to prevent fatal intestinal damage and sepsis during acute schistosomiasis; however, AAMs can also produce a variety of factors to recruit and activate fibroblasts, which contribute to the development of fibrosis [2, 12]. Depletion of macrophages/ monocytes attenuates liver and lung granuloma formation and tissue fibrosis after *Schistosoma* infection [3, 13]. Thus, preventing excessive monocyte infiltration is important for tissue repair and host survival in chronic schistosomiasis. Nevertheless, despite the clear and well-documented roles of monocytes and macrophages in schistosomiasis, little is known about the mechanisms underlying regulation of monocyte infiltration.

Infection with *Schistosoma* induces IL-10–producing B cells, a relatively new member in the network of regulatory immune cells [14, 15]. *Schistosoma mansoni*– infected B cell–deficient μMT mice show more extensive hepatic granulomas and fibrosis than WT mice [16-18], but the mechanisms underpinning this difference are unclear. In mice, two major populations of B cells exist: B1 cells and B2 cells. On the basis of cluster of differentiation (CD)5 expression, B1 cells can be further subdivided into B1a (CD5+) and B1b (CD5−) subsets [19-21]. The B1 cells reside mainly in the peritoneal and pleural cavities, with low frequencies (<5%) in the spleen. The B1a cells spontaneously secrete natural IgM antibodies, which bind self-antigens, bacterial cell wall components, or viruses [22, 23]. The B1a cells also spontaneously secrete IL-10, which regulates acute and chronic inflammatory diseases [19]. In the present study, we investigated the cross talk between B1a cells and monocytes to understand their roles in the pathogenesis of schistosomiasis. By using a murine model of *Schistosoma japonicum* infection, we demonstrated that B1a cells suppress granulomatous inflammation and liver fibrosis by regulating Ly6C^hi^ monocyte infiltration. We also found that IL-10 was required for B1a cells to downregulate the expression of chemokines and cytokines that attract monocytes.

## Results

### B cells protect against *S. japonicum*–induced liver pathology

To assess the role of B cells in the liver pathology associated with schistosomiasis, we infected B cell–deficient (μMT) mice and WT mice with *S. japonicum* and harvested samples at the indicated times (Fig 1A). We found that the sizes of the hepatic granulomas after infection in μMT mice were greater than those in WT mice (Fig 1B and D). Liver fibrosis was measured using picrosirius red staining and hydroxyproline levels. The results showed that both the proportion of the collagen area and the hepatic hydroxyproline levels in μMT mice 8 weeks and 10 weeks after infection were increased compared with those in WT mice (Fig 1C, E and F), indicating that μMT mice exhibited increased hepatic fibrosis. In addition, serum alanine transaminase (ALT) levels were significantly higher in μMT mice 6 weeks after infection (Fig 1G), suggesting that liver injury is more severe in μMT mice than in WT mice. The more severe liver pathology in μMT mice was not due to an increased burden of infection because the numbers of eggs observed in the liver samples did not differ significantly between μMT mice and WT mice (S1 Fig). Together, these data reveal an important role for B cells in attenuating *S. japonicum* egg-induced granuloma formation, hepatic injury, and hepatic fibrosis.

**Fig. 1.**
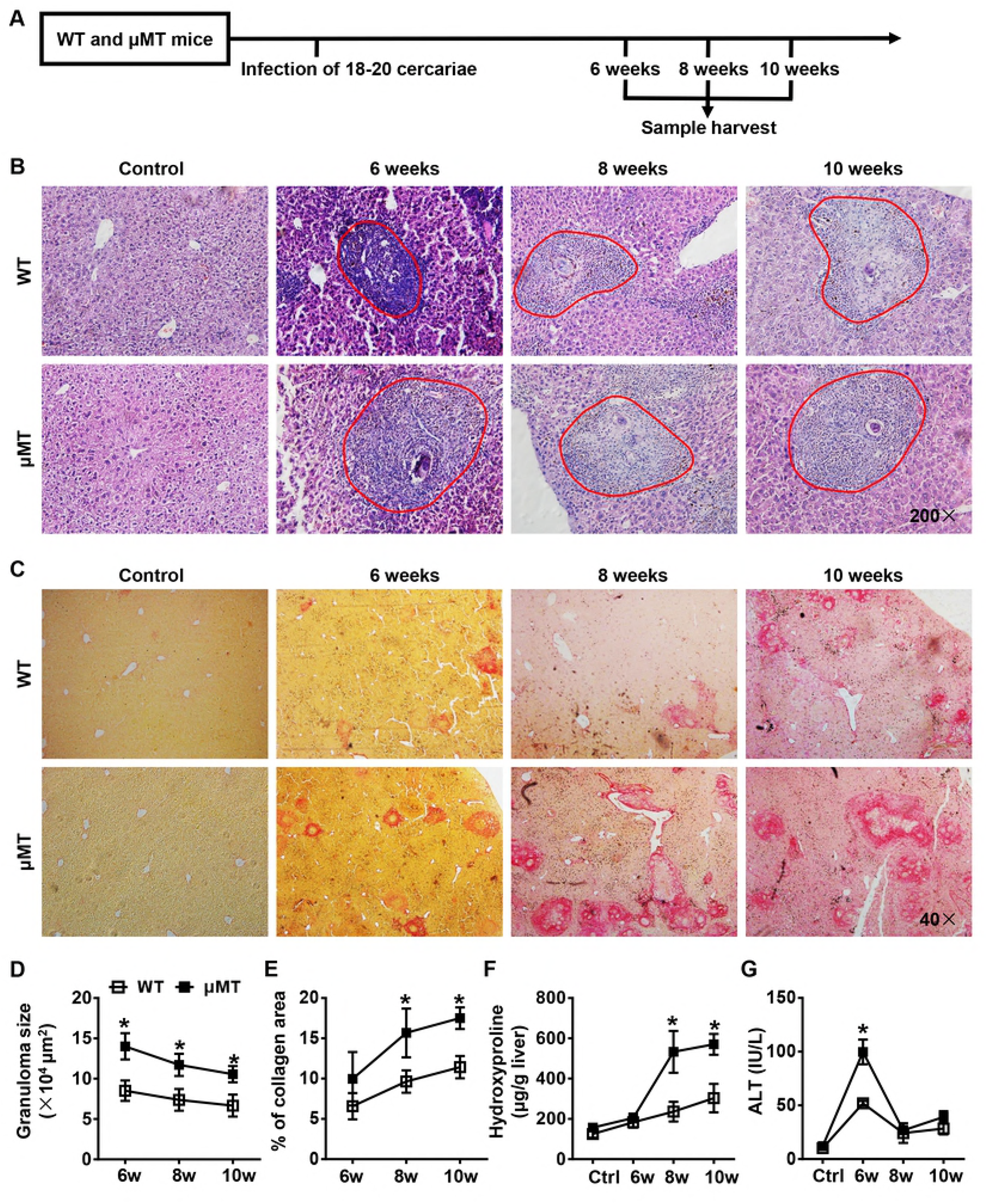
Mice lacking B cells exhibit more severe liver pathology than wild-type mice after *S. japonicum* infection. (**A**) Schematic representation of the model of *S. japonicum* infection. WT mice and μMT mice were infected with 18–20 cercariae of *S. japonicum*, and liver samples from these mice were harvested at the times indicated after infection. Comparisons were made with uninfected control mice. (**B, C**) Representative graphs of hematoxylin-eosin staining (**B**) and picrosirius red staining (**C**) of liver specimens. (B) All images were taken at 200 × magnification. The red outlined areas indicate granulomas. (C) All images were taken at 40 × magnification. (**D-G**) Statistical analysis of granuloma sizes (**D**), proportion of collagen areas (**E**), amount of hepatic hydroxyproline (**F**) and levels of serum alanine transaminase (ALT) (**G**). Data represent mean ± SD; n = 5–7 per time point from two independent experiments. \**p* < 0.05, versus WT mice, two-tailed, unpaired Student’s *t* test.

### B cells regulate the recruitment of monocytes in the liver

To further analyze the cellular components in the granulomas, we isolated hepatic leukocytes and used flow cytometry to detect cell phenotypes. The results showed that compared with those of WT mice the numbers of leukocytes (CD45^+^) and total macrophages (CD11b^+^F4/80^+^) were markedly increased in μMT mice during all stages of infection. The numbers of neutrophils (CD11b^+^Ly6G^+^), and of natural killer T (NKT) cells (CD3^+^NK1.1^+^), were modestly but significantly increased in μMT mice 8 weeks after infection, while the numbers of T cells (CD3^+^NK1.1^−^) and NK cells (CD3^−^NK1.1^+^) showed no significant changes between WT mice and μMT mice (Fig 2).

**Fig. 2.**
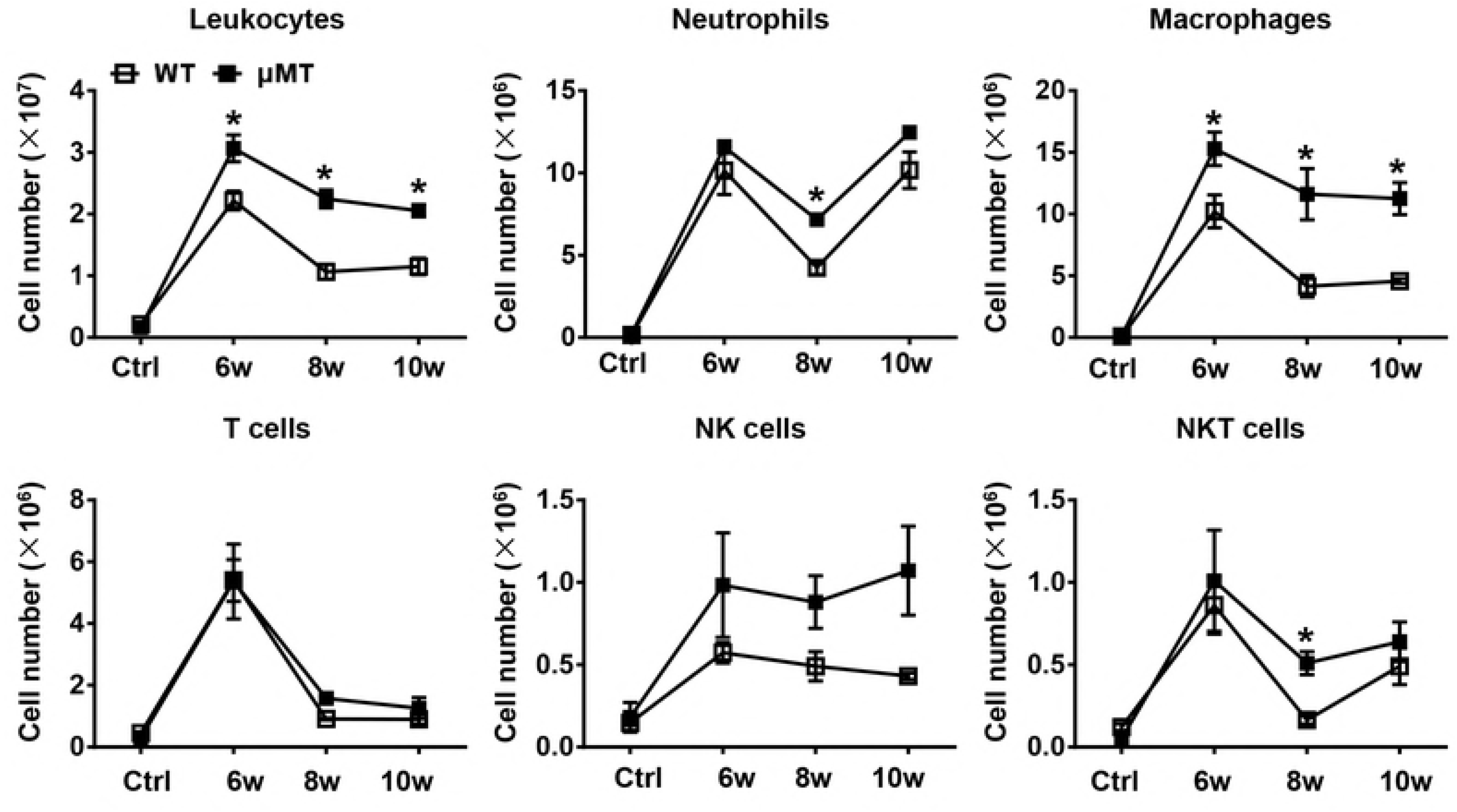
B cell deficiency results in increased number of macrophages. The infiltration of hepatic leukocytes (CD45^+^), macrophages (CD11b^+^F4/80^+^), neutrophils (CD11b^+^Ly6G^+^), T cells (CD3^+^NK1.1^−^), NK cells (CD3−NK1.1^+^), and NKT cells (CD3^+^NK1.1^+^) after infection were quantified by flow cytometric analysis. Controls (Ctrl) were uninfected mice. Data represent mean ± SD; n = 3–5 per time point from three independent experiment. \**p* < 0.05, versus WT mice, two-tailed, unpaired Student’s *t* test.

Hepatic macrophages consist of distinct populations termed resident Kupffer cells and monocyte-derived macrophages (MoMFs). The MoMFs can be further identified as two distinct subsets: proinflammatory Ly6C^hi^ MoMFs and restorative Ly6C^lo^ MoMFs [24]. In the present study, we found that a larger number of proinflammatory Ly6C^hi^ monocytes infiltrated the liver in μMT mice than in WT mice 6 weeks after infection. By contrast, the numbers of Kupffer cells and Ly6C^lo^ MoMFs were similar in μMT mice and WT mice (Fig 3).

**Fig. 3.**
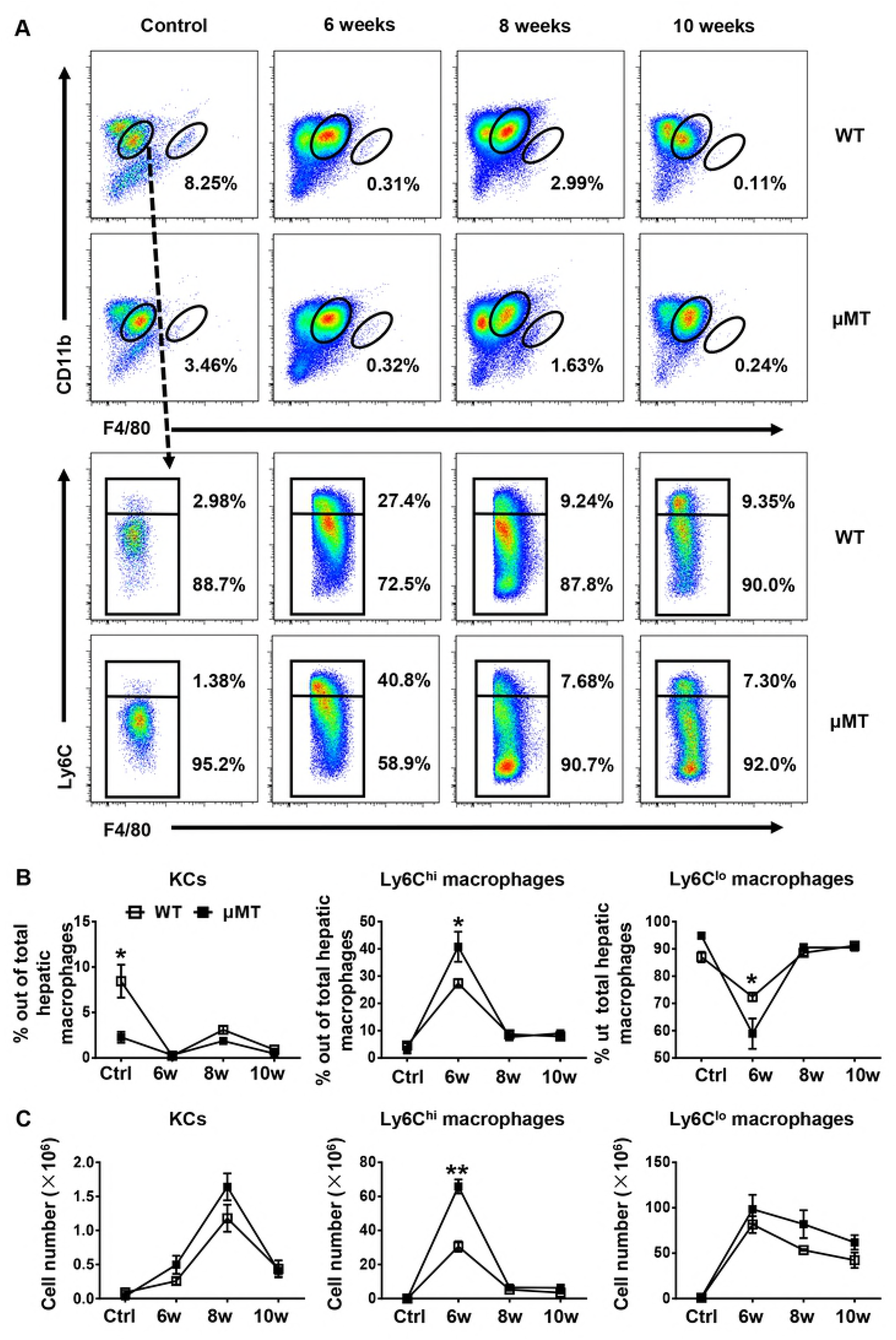
Ly6C^hi^ MoMFs are significantly increased in μMT mice. Flow cytometric analysis of macrophage subsets in WT mice and μMT mice after *S. japonicum* infection. (**A**) Representative fluorescence-activated cell sorting plots are shown for the indicated times after *S. japonicum* infection. Upper panels are pre-gated on CD45^+^ and live cells. Bottom panels are pre-gated on CD45^+^CD11bhiF4/80lo cells. (**B**) Graphical summary showing percentage of Kupffer cell (KCs) (CD11bloF4/80hi), Ly6C^hi^ MoMF (CD11bhiF4/80loLy6C^hi^), and Ly6C^lo^ MoMF (CD11bhiF4/80loLy6C^lo^) subsets out of total hepatic macrophags. (C) Absolute numbers of KCs, Ly6C^hi^ MoMFs, and Ly6C^lo^ MoMFs in WT mice and μMT mice. Data represent mean ± SD; n = 8–10 per time point from three independent experiments. \**p* < 0.05, \*\**p* < 0.01, versus WT mice, two-tailed, unpaired Student’s *t* test.

We hypothesized that the increased macrophage number in the liver of μMT mice after infection reflects either enhanced monocyte production or increased monocyte recruitment. Because Ly6C^hi^ monocytes are derived from the bone marrow and circulate in the blood [25], we analyzed Ly6C^hi^ monocytes in the peripheral blood. We found no significant difference in the number of circulating Ly6C^hi^ monocytes after infection, despite somewhat higher counts in uninfected naïve μMT mice, which suggests that monocyte production does not account for the increased monocyte number in the liver of μMT mice (S2 Fig). To evaluate recruitment, we used quantitative polymerase chain reaction (PCR) assays to examine the gene expression levels of chemokines that attract monocytes in the liver samples of WT mice and μMT mice 6 weeks after infection. The expression levels of *Ccl1, Ccl2, Ccl3, Ccl4*, and *Ccl5* were higher in the liver of μMT mice than in WT mice (Fig 4A). We also detected the protein levels of some key chemokines. The protein levels of CCL2, CCL3, CCL4, and CCL5 in the liver of μMT mice were significantly increased compared with those of WT mice (Fig 4B). These data suggest that mobilization and recruitment, rather than production, accounted for differences in monocyte infiltration. Therefore, B cells limit monocyte influx by suppressing the expression of chemokines during the acute stage of *S. japonicum* infection.

**Fig. 4.**
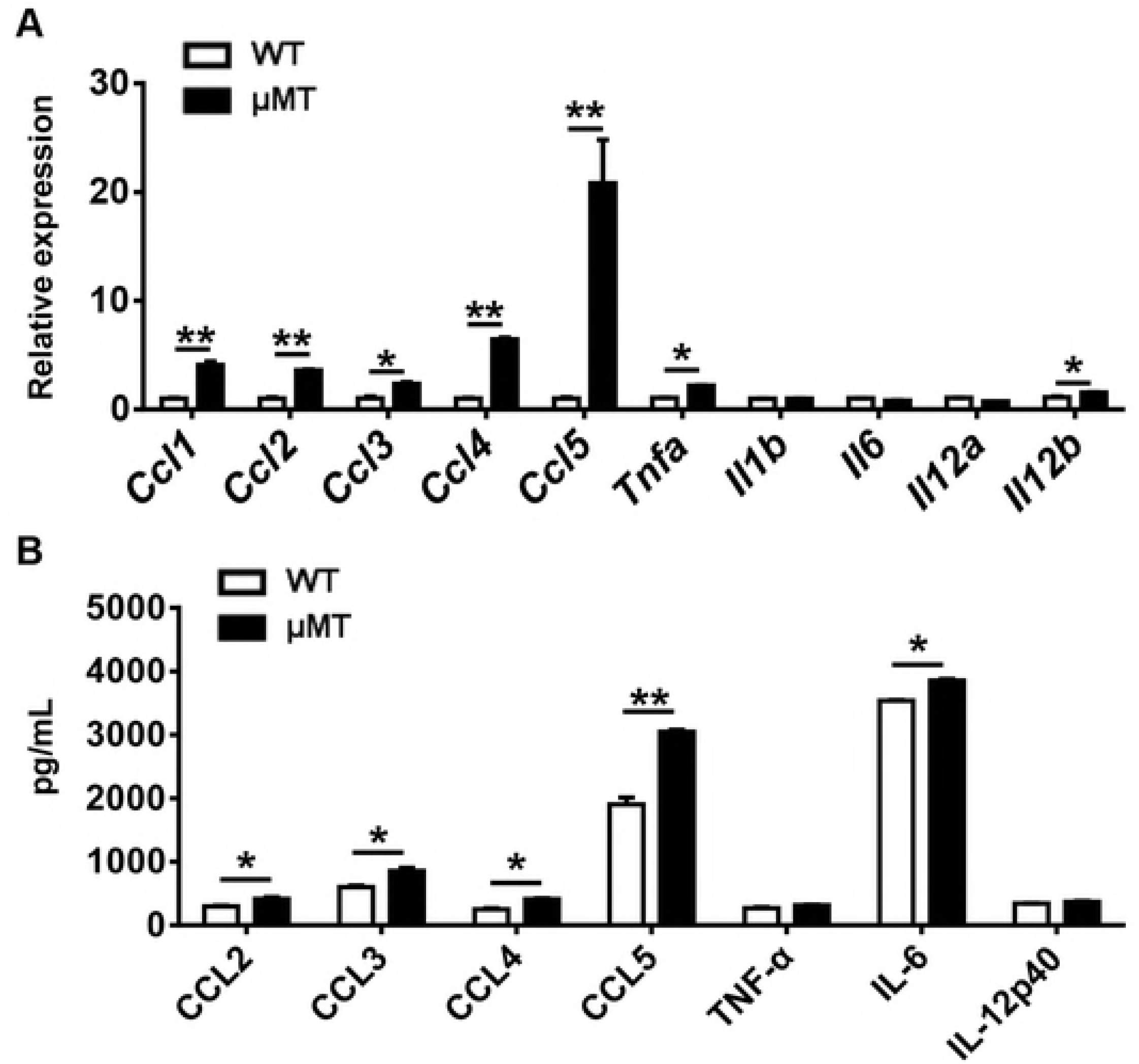
Inflammatory cytokines and chemokines are increased in the liver of μMT mice after *S. japonicum* infection. (**A**) Relative gene expression of chemokines (*Ccl1, Ccl2, Ccl3, Ccl4,* and *Ccl5*) and inflammatory cytokines (*Tnfa, Il1b, Il6, Il10, Il12a* and *Il12b*) in the livers of WT mice and μMT mice 6 weeks after infection. Data are from three independent experiments. (**B**) Protein levels of CCL2, CCL3, CCL4, CCL5, TNF-α, IL-6, and IL-12p40 in liver 6 weeks after the infection (n = 5–7 per group from two independent experiments). Data represent mean ± SD. \**p* < 0.05, \*\**p* < 0.01, two-tailed, unpaired Student’s *t* test.

Ly6C^hi^ MoMFs preferentially express inflammatory cytokines [26]. Thus, we compared the expression of selected cytokines in the livers of μMT mice and WT mice after *S. japonicum* infection. The gene expression levels of *Tnfa* and *Il12b* and the protein levels of IL-6 in the livers of μMT mice were significantly higher than those in WT mice (Fig 4). In addition, serum protein levels of CCL3 and CCL5 in μMT mice were lower than those in WT mice (S3 Fig).

### B1a cells migrate from the peritoneal cavity (PC) to the liver after infection

To investigate the role and mechanism of B cell action in this murine model, we first determined the number of total B cells in the liver during infection. We found that the B cell number was significantly increased 6 weeks after infection in WT mice (Fig 5A). Further analysis of hepatic B cell subsets showed that both the percentage and number of hepatic B1a cells were markedly increased 6 weeks after infection, and the numbers of B1b cells and B2 cells were also increased (Fig 5B, 5C, and S4 Fig). In addition, we found that both the percentage and number of PC B1a cells were markedly decreased 6 weeks after infection, whereas the percentage and number of PC B2 cells were increased (Fig 5D and E). These data suggest that the increased B1 cells in the liver after infection were recruited from the PC. To provide further support for this finding, we conducted adoptive cell transfer experiments. B cell–deficient μMT mice were infected with *S. japonicum* and then were intraperitoneally injected with uninfected WT mice–derived PC B cells or phosphate-buffered saline (PBS) 4 weeks after infection. Samples were harvested 6 weeks after infection (Fig 6A). The purity of WT mice–derived PC B cells was more than 95% (Fig 6B). After the cell transfer, B cell subsets could be detected in the PC and liver of μMT mice. Compared with those in donor WT mice, the percentage of B1a cells in the PC was lower in the recipient μMT mice, whereas the percentage of B1a cells in the liver was higher, which was consist with our observations in the infected WT mice (Fig 6C-F).

**Fig. 5.**
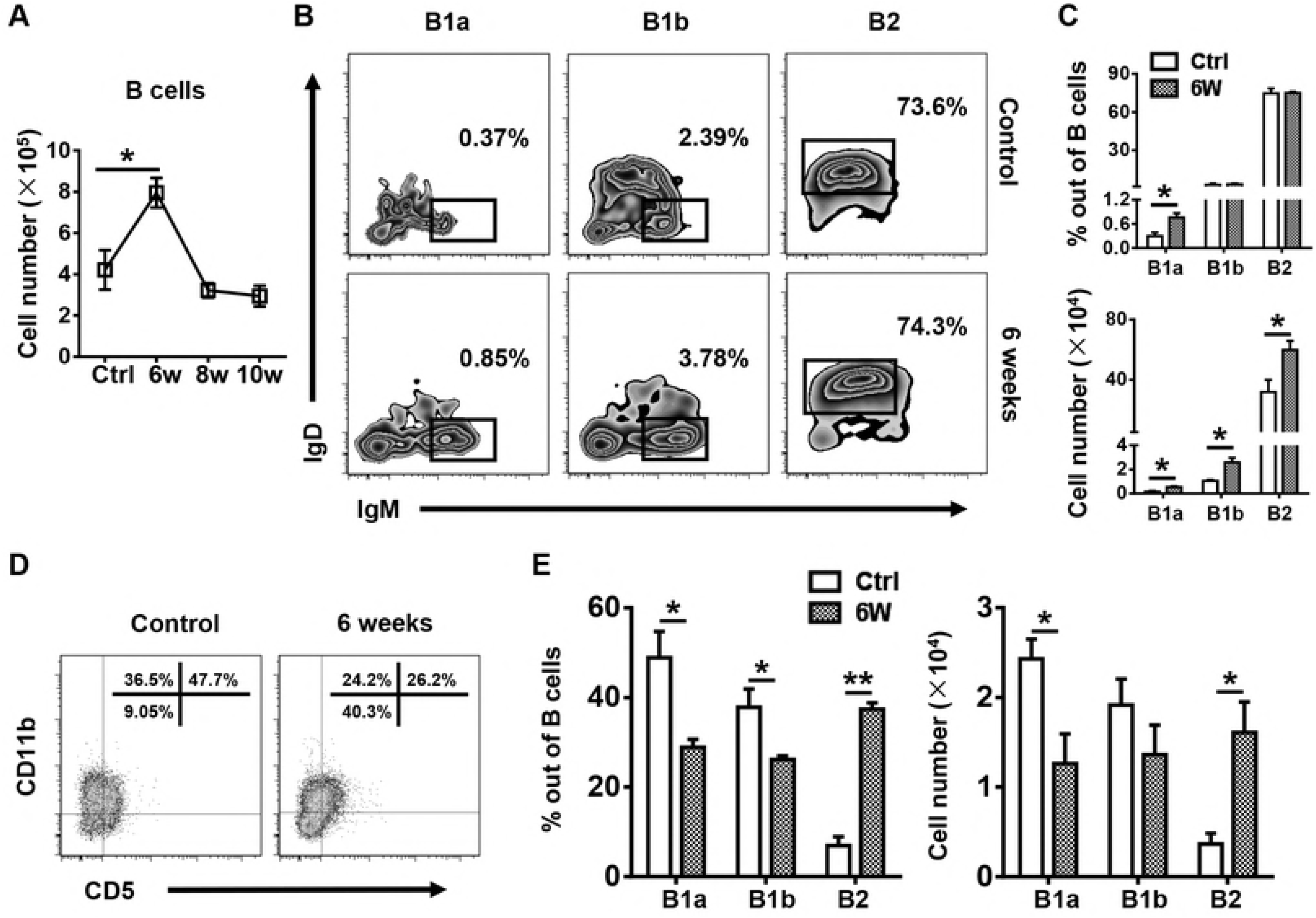
Hepatic B1a cells increase whereas peritoneal cavity (PC) B1a cells decrease after *S. japonicum* infection. (**A**) Number of B cells after infection in WT mice. (**B**) Representative flow cytometry plots of hepatic B1a, B1b, and B2 cells in WT mice 6 weeks after infection. (**C**) Graphical summary showing the percentage of B1a, B1b, and B2 cells out of total B cells (top panel) and the number of indicated subsets (bottom panel) in the livers of WT mice without infection (Ctrl) and 6 weeks after infection. (**D**) Representative flow cytometry plots of PC B1a, B1b, and B2 cells in WT mice without infection and 6 weeks after infection. (**E**) The percentage of B1a, B1b, and B2 cells out of total B cells (left panel) and number of indicated subsets (right panel) in the PC of WT mice without infection and 6 weeks after infection. Data represent mean ± SD; n = 5–7 per group from two independent experiments. \**p* < 0.05, \*\**p* < 0.01, two-tailed, unpaired Student’s *t* test.

**Fig. 6.**
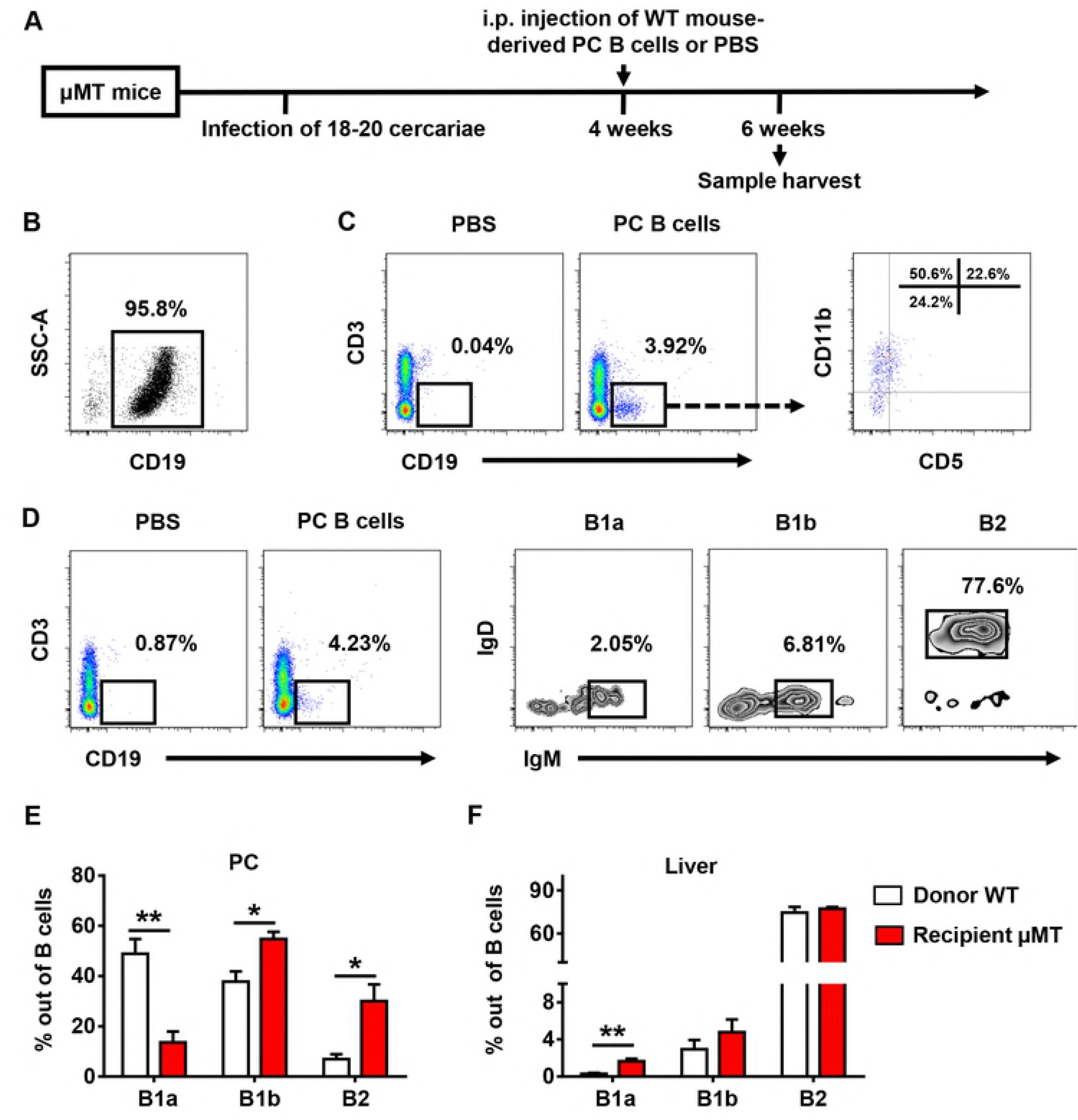
Transferred peritoneal cavity (PC) B1a cells preferentially accumulate in the livers of receiving μMT mice. (**A**) Schematic representation of the PC B cell transfer. (**B**) Purity of PC B cells from WT mice after sorting. (**C, D**) Flow cytometric analysis of PC (**C**) and liver (**D**) B cell subsets after transfer in μMT mice. (**E, F**) The frequencies of B1a, B1b, and B2 cells in PC (**E**) and liver (**F**) of donor WT mice and recipient μMT mice. Data represent mean ± SD; n = 5–7 per group from two independent experiments. \**p* < 0.05, \*\**p* < 0.01, two-tailed, unpaired Student’s *t*-test.

### IL-10 is indispensable for B cell protection against *S. japonicum* infection–induced liver pathology

The regulatory function of B cells is mediated mainly by their secretion of IL-10 [21, 27-30]. To determine whether B cells protect against *S. japonicum* infection– induced liver pathology via IL-10, we first examined IL-10 expression in the liver and B cells. The hepatic IL-10 protein levels in μMT mice were significantly lower than those of WT mice 6 weeks after infection (Fig 7B), suggesting that B cells contribute to IL-10 production in the liver after infection. Our data also showed that IL-10 expression levels in B cells were increased after infection (Fig 7C, D), especially in hepatic B1a cells (Fig 7E).

**Fig. 7.**
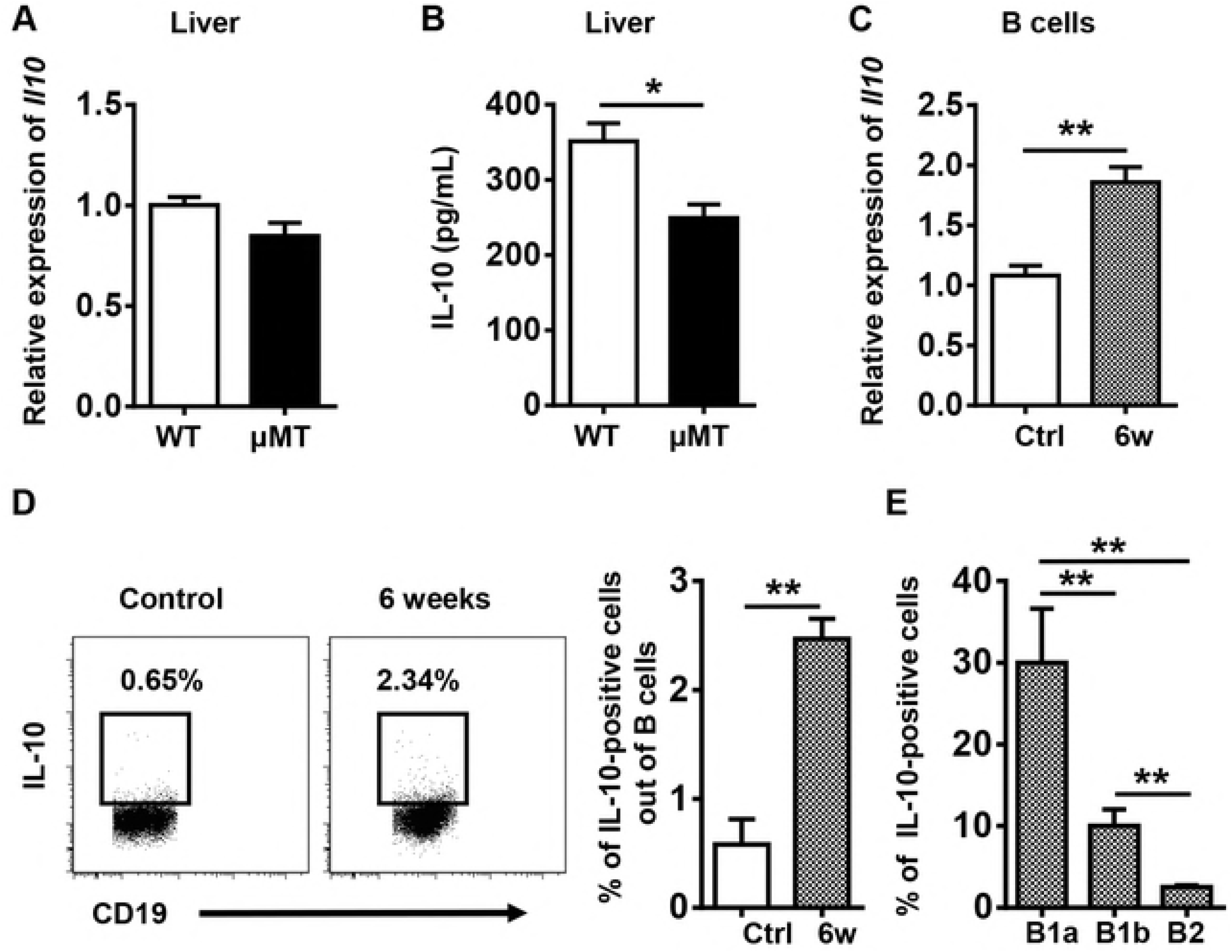
IL-10 expression is increased in the liver and in B cells after *S. japonicum* infection. (**A**) Relative *Il10* gene expression in the livers of WT mice and μMT mice 6 weeks after infection; data are from three independent experiments. (**B**) IL-10 protein levels in the livers of WT mice and μMT mice 6 weeks after infection. (**C**) Quantitative PCR analysis of *Il10* in sorted hepatic B cells; data are from three independent experiments. (**D**) The frequency of IL-10–positive cells out of total B cells in the livers of WT mice was examined by flow cytometry. (**E**) Graphical summary showing percentage of IL-10–positive cells out of indicated cell subsets in WT mice 6 weeks after infection. Data represent mean ± SD; n = 5–7 per group from two independent experiments. \**p* < 0.05, \*\**p* < 0.01, two-tailed, unpaired Student’s *t* test.

IL-10 plays a protective immunomodulatory role during schistosomiasis [16]. To assess whether IL-10 is involved in the suppressive effect of B cells on monocyte infiltration after *S. japonicum* infection, we adoptively transferred PC B cells from WT or *Il10*^−/−^ mice into *S. japonicum*–infected μMT mice. As expected, the μMT mice that received WT B cells showed decreases in the granuloma sizes, ALT levels, numbers of Kupffer cells and Ly6C^hi^/Ly6C^lo^ MoMFs, and the expression levels of hepatic chemokines and inflammatory cytokines compared with those in control mice receiving PBS (Fig 8A-G). However, when μMT mice received the IL-10–deficient B cells, the granuloma sizes, numbers of Kupffer cells and Ly6C^hi^/Ly6C^lo^ MoMFs, and expression levels of chemokines and inflammatory cytokines in the liver were not reduced compared with those in controls (Fig 8A, B, and E-G). Only the ALT levels in μMT mice receiving the transfer of IL-10–deficient B cells were deceased (Fig 8C). Collectively, these results provide evidence that the B1a regulatory subset of B cells suppress monocyte recruitment by producing IL-10 to thereby attenuate *S. japonicum* egg–induced liver pathology.

**Fig. 8.**
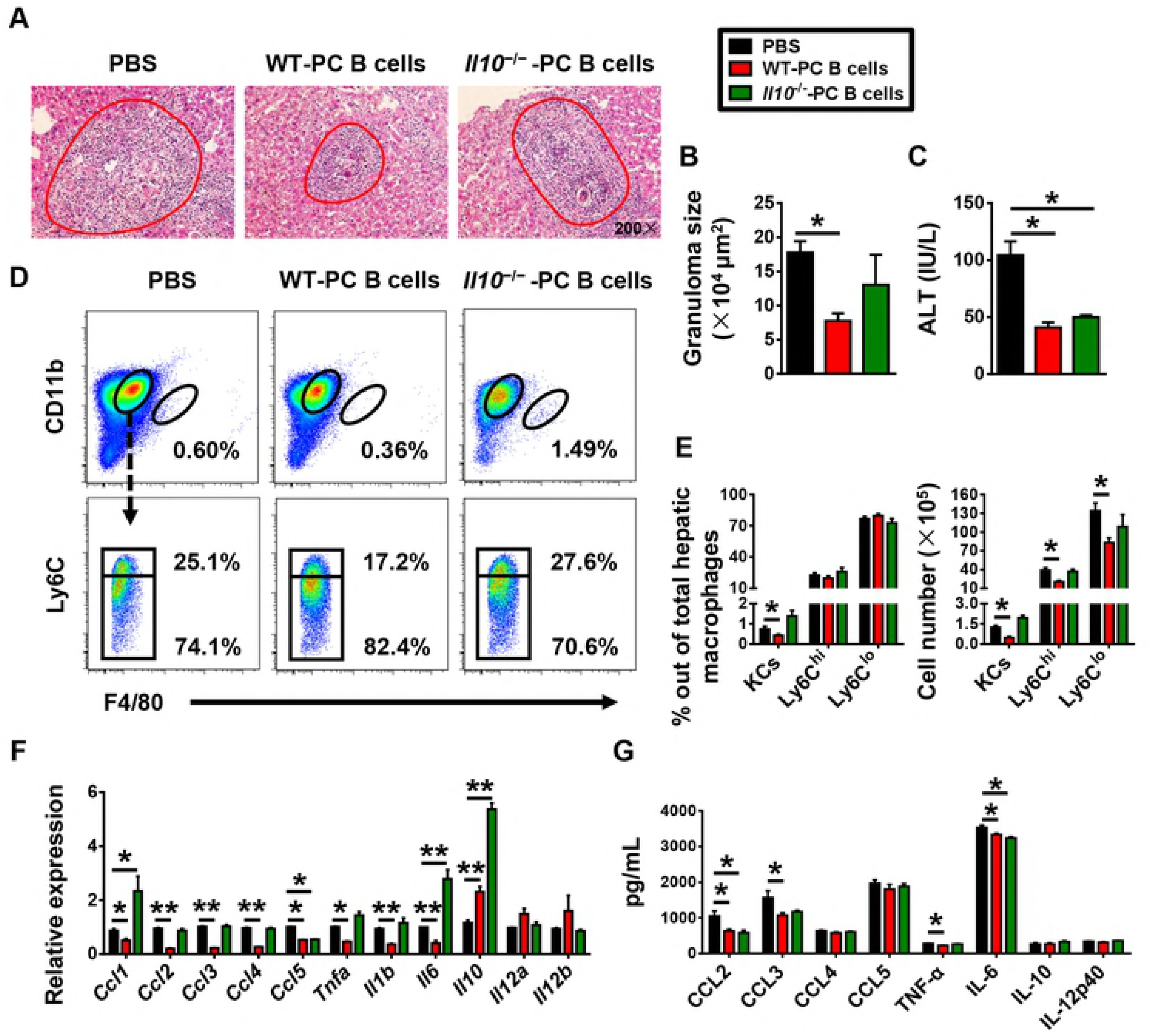
Adoptive transfer of WT PC B cells, but not IL10^-/-^ PC B cells, attenuates *S. japonicum*–induced liver pathology in μMT mice. The μMT mice were infected with 18–20 cercariae of *S. japonicum*. The adoptive transfer of B cells (1 × 106 cells) purified from the PC of WT or IL-10^−/−^ mice into μMT mice was performed 4 weeks after infection. Mice were sacrificed 6 weeks after infection. (**A**) Representative images of hematoxylin-eosin stained liver tissues. (**B**) Statistical analysis of hepatic granuloma sizes and (**C**) serum ALT levels. (**D**) Representative flow cytometry plots of hepatic macrophage subsets after cell transfer. (**E**) Left panel: graphical summary showing percentage of indicated cell subsets out of total hepatic macrophages in infected μMT mice after cell transfer. Right panel: number of indicated cell subsets in the liver of infected μMT mice after cell transfer. (**F**) Quantitative PCR analysis of chemokine and inflammatory cytokine gene expression levels in the liver; data are from three independent experiments. (**G**) Hepatic chemokine and inflammatory cytokine protein expression levels were examined by cytometric bead assay and enzyme-linked immunosorbent assay. Data represent mean ± SD; n = 5–7 per group from two independent experiments. \**p* < 0.05, \*\**p* < 0.01, two-tailed, unpaired Student’s *t* test.

## Discussion

The roles of B cells in liver fibrosis remain obscure. In carbon tetrachloride (CCl_4_)- induced liver fibrosis, B cells are required for the fibrotic processes. In the CCl_4_ model, B cells serve to amplify liver fibrosis though the production of proinflammatory cytokines and chemokines [31, 32]. However, the observed mechanisms in the CCl_4_ model are not applicable to infectious liver fibrosis. The B cell–deficient mouse displays an increased hepatic fibrosis after *Schistosoma mansoni* infection, suggesting that B cells serve a protective role in infection-induced liver fibrosis. The mechanisms underlying B cell suppression of *Schistosoma*-induced liver fibrosis had been previously unknown. However, in the present study using an *S. japonicum*–infected murine model, we found that B1 cells protected against *S. japonicum* infection–induced liver pathology by controlling liver infiltration of monocytes. In agreement with the reports using the *S. mansoni* infection model [16, 17], we observed markedly exacerbated hepatic granuloma formation, liver injury, and fibrosis in *S. japonicum*– infected B cell–deficient (μMT) mice. The B1a cells trafficked from the peritoneal cavity to the liver following infection induction. The increased B1a cells in the liver suppressed the production of chemokines, which attract monocytes, and thus controlled the recruitment of monocytes. The B1a cells played their regulatory roles via producing IL-10 (Fig 9).

**Fig. 9.**
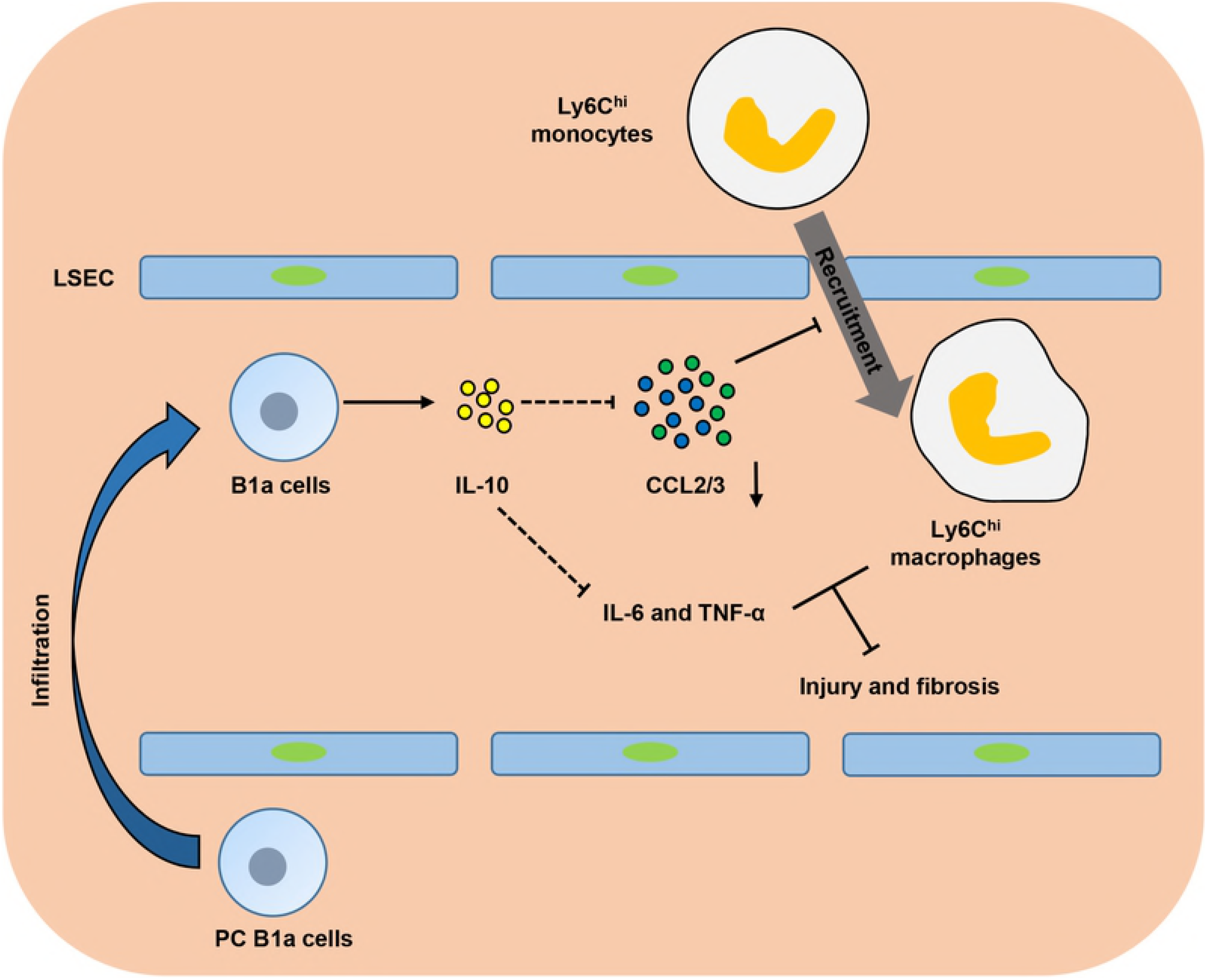
Model describing B1a cell suppression of *S. japonicum*–induced liver pathology. After *S. japonicum* infection, peritoneal cavity (PC) B1a cells infiltrate the liver. These increased numbers of hepatic B1a cells secrete IL-10, which downregulates the expression of hepatic CCL2 and CCL3 to inhibit excessive hepatic infiltration of Ly6C^hi^ monocytes. Thus, B1a cells alleviate liver early inflammation and late fibrosis. LSEC, liver sinusoidal endothelial cells.

Infiltrating Ly6C^hi^ monocytes may act as a double-edged sword in liver damage. These cells express a substantial number of inflammatory cytokines and chemokines and promote liver inflammation, injury, and fibrosis in the initiation and progression of various types of liver injury, including acute viral injection, hepatotoxicity following CCl_4_ treatment, or ischemia–reperfusion damage [24, 33, 34]. We hypothesized that recruited Ly6C^hi^ monocytes also contribute to the initial liver damage and development of fibrosis after *S. japonicum* infection. As expected, *S. japonicum*–infected μMT mice with increased Ly6C^hi^ monocyte infiltration had higher levels of ALT and liver fibrosis than WT mice (Fig 1 and 3). When liver injury ceases, inflammatory Ly6C^hi^ monocytes mature into Ly6C^lo^ restorative macrophages, which display increased expression of anti-inflammatory cytokines, regenerative growth factors, and matrix degrading metalloproteinase [9, 25]. During chronic *S. mansoni* infection, Ly6C^hi^ monocytes become AAMs in granulomas through a Ly6C^lo^ state [3, 4]. The arginase 1–expressing AAM population suppresses Th2 cytokine-driven inflammation and fibrosis in schistosomiasis [35, 36]. Thus, it is crucial to regulate monocyte recruitment and homeostasis in the liver. Our results showed that compared with WT mice, μMT mice had increased hepatic Ly6C^hi^ MoMFs and chemokines attracting Ly6C^hi^ monocytes after *S. japonicum* infection (Fig 3 and 4), which suggests that B cells play critical roles in controlling monocyte infiltration after *S. japonicum* infection through negative regulation of chemokines.

Schistosoma infection induces IL-10–producing B cells, which are termed regulatory B cells or B10 cells [27, 30]. Currently, there are no phenotypic, transcription factors, or lineage markers that are unique to B10 cells, and B10 cells mostly overlap with B1 cells [19, 21, 28]. B1 cells can secrete IL-10 to mediate the negative regulation of inflammation, including restricting the production of proinflammatory cytokines, downregulating the expression of major histocompatibility complex class II [37], and maintaining the suppressive function of regulatory T cells [38]. In homeostatic conditions, B1a cells are localized mainly in the PC, and they are the major population of B cells in this compartment. In response to pathogens, serosal B1a cells in body cavities migrate to neighboring lymphoid sites or tissues [19, 39]. In the present study, we found that B1 cells, especially B1a cells, migrated from the PC to the liver after *S. japonicum* infection, which was shown by the increased percentage and number of B1a cells in the liver and their concurrent decrease in the PC after infection (Fig 5). In addition, *S. japonicum*–infected μMT mice receiving the adoptive transfer of PC B cells purified from WT mice also showed a higher percentage of B1a cells in the liver and a lower percentage of B1a cells in the PC than the donor mice (Fig 6). These data suggest that B1a cells are the major population of B cells that regulate monocyte infiltration after *S. japonicum* infection.

IL-10 is involved in the immunoregulatory role of B1a cells [19]. It has been reported that IL-10 plays an antifibrotic role via inhibiting the proliferation and collagen synthesis of myofibroblasts [40]. Recently, a connection between IL-10 and inflammatory chemokines has been suggested in renal fibrosis and nerve injury. After the onset of unilateral ureteral obstruction, IL-10 knockout mice show increased infiltration of inflammatory cells and cytokines, including monocyte chemoattractant protein-1, TNF-α, IL-6, IL-8, RANTES, or macrophage colony-stimulating factor, in the kidney compared with WT controls [41]. After peripheral nerve injury, IL-10 plays a role in controlling the early influx and the later efflux of macrophages out of the nerve via downregulating expression of proinflammatory chemokines and cytokines [42]. In the present study, we observed an increased expression of IL-10 in B cells. Our flow cytometric analysis indicated that B1a cells are the major population of B cells expressing IL-10 in the *S. japonicum*–infected liver (Fig 7). We also found that in the absence of IL-10, the transferred PC B cells were unable to downregulate granuloma inflammation, recruitment of monocytes, or the expression of a number of proinflammatory chemokines and cytokines in the infected μMT mice (Fig 8). These data suggest that after *S. japonicum* infection, B cells control the recruitment of monocytes and the expression of proinflammatory chemokines and cytokines via IL-10 production.

The cross talk between B cells and monocytes observed in our study appears to be opposite to that observed in CCl_4_-induced fibrosis [31]. The difference may be that different liver microenvironments induce different B cell subsets in these two models. In the CCl_4_ model, the increased B cells in the liver produce IgG and express CD138, which may be a B2 subset. The B cells in the CCl_4_ model secrete proinflammatory cytokines and chemokines, hence, they recruit dendritic cells and Ly6C^++^ monocytes. In the present model, the B1a cells were the most substantially increased B cell subset in the *S. japonicum*–infected liver. These B1a cells produced IL-10, which led to the suppressed recruitment of Ly6C^hi^ monocytes.

In conclusion, our data indicated that PC B1a cells infiltrate the liver when it is damaged through *S. japonicum* infection. These B1a cells secrete IL-10, which inhibits expression of CCL2, CCL4, and CCL5 to limit excessive liver infiltration of Ly6C^hi^ monocytes and thereby alleviate early inflammation and later liver fibrosis.

## Materials and methods

### Ethics statement

All animal experiments were approved by the Institutional Animal Care and Use Committee at Anhui Medical University (the approved number is LLSC20150279) and conformed to the guidelines outlined in the Guide for the Care and Use of Laboratory Animals.

### Mice and parasites

The 8- to 10-week-old male C57BL/6 WT mice and B cell–deficient (μMT) mice on a C57BL/6 background were purchased from the Nanjing Biomedical Research Institute of Nanjing University (Nanjing, China). The *Il10*^−/−^ mice on a C57BL/6 background were provided by Professor Zhigang Tian (University of Science and Technology of China). All mice were kept under temperature- and humidity-controlled specific-pathogen-free conditions. For infection, mice were anesthetized and percutaneously exposed to 18–20 cercariae of *S. japonicum* (a Chinese mainland strain) that were obtained from infected *Oncomelania hupensis* snails. At the indicated times, mice were euthanized and tissue samples were harvested for later experiments.

### Egg counts

A portion of the liver tissue was digested in 10% potassium hydroxide at 37°C for 3 h. The eggs in aliquots of the suspensions were counted under a microscope.

### Cell isolation

For isolation of peripheral leukocytes, blood samples were incubated with ACK (Ammonium-Chloride-Potassium) Lysis Buffer (GibcoTM) on ice for 10 min to remove red blood cells. After being neutralized and washed, the pellets were resuspended with PBS.

For isolation of PC cells, the outer layer of the peritoneum was opened. A needle was inserted into the inner layer of the peritoneum to avoid puncture of organs. Ice-cold PBS (5 mL) containing 2% bovine serum albumin was injected into the PC and the PC was washed repeatedly. The collected cell suspension was centrifuged at 500 × *g* for 10 min, and the pellets were resuspended with PBS.

For isolation of hepatic leukocytes, liver samples were cut and incubated in Dulbecco’s modified Eagle’s medium (DMEM) containing 0.05% collagenase IV (Sigma), 0.002% DNase I (Sangon Biotech Co. Ltd.) and 10 mM HEPES at 37°C for 40 min. The digested liver tissue was passed through nylon mesh (74 μm), and the enzymes were inactivated by adding additional DMEM. After centrifugation at 50 × *g* twice for 2 min to remove the hepatocyte pellet, the supernatant was centrifuged at 500 × *g* for 10 min. The pellets were resuspended in 40% Percoll (GE Healthcare) and centrifuged at 1260 × *g* for 20 min. The resulting pellets were incubated for 5 min on ice with ACK Lysis Buffer and resuspended with PBS.

All single-cell suspensions were washed and counted. Cell viability was confirmed using the standard trypan blue exclusion method.

Liver and PC CD19^+^ B cells were sorted by PE-CD19 and anti-PE microbeads using a magnetic affinity cell sorting (MACS) system (Miltenyi Biotec), and the purity was >90%.

### Adoptive transfer of PC B cells

MACS-sorted uninfected WT or *Il10*^−/−^ mouse–derived PC B cells (2 × 10^6^) or PBS was intraperitoneally injected into each recipient μMT mouse 4 weeks post infection.

### Flow cytometry

The following fluorochrome-conjugated monoclonal antibodies were used in this study: antimouse CD3, CD5, CD11b, CD19, CD23, CD45, CD115, F4/80, Ly6C, Ly6G, NK1.1, IgM, IgD (all from BioLegend, San Diego, CA), IL-10 (BD Pharmingen). The cells (1×10^6^) were blocked with FcR blocker (BD Pharmingen) and then incubated with monoclonal antibodies to surface antigens.

To detect the secretion of IL-10, cells (1 × 10^6^) were stimulated with phorbol myristate acetate (30 ng/mL), ionomycin (1 μg/mL) (Sigma, St. Louis, Mo.), monensin (5 μg/mL) (Sigma, St. Louis, Mo.), and lipopolysaccharide (10 μg/mL) (Sigma, St. Louis, Mo.) for 4 h. Cells were incubated with monoclonal antibodies to surface antigens and then fixed and permeabilized using a Transcription Factor Staining Buffer Set (eBioscience, San Diego, CA). The cells were then incubated with antibodies to IL-10. All samples were analyzed using flow cytometry (FACSVerse system, BD Biosciences) with FlowJo (version 7.6.1) software.

### RNA isolation and quantitative PCR

Total hepatic RNA was isolated from frozen liver tissue using Trizol (Invitrogen). The MACS-sorted hepatic B cells were resuspended in Trizol, and the RNA was isolated according to the manufacturer’s instructions. First strand cDNA was synthesized from ≤500 ng of RNA using a PrimeScript RT reagent kit (TaKaRa). Quantitative PCR was performed with a StepOnePlus Real-Time PCR System (Applied Biosystems, Foster City, CA) using SYBR Premix Ex Taq II (TaKaRa). The expression levels of target genes were normalized to the housekeeping gene *Actb*. Relative expression was calculated by the 2−ΔΔCt method. The primers used are given in Table 1.

**Table 1.**
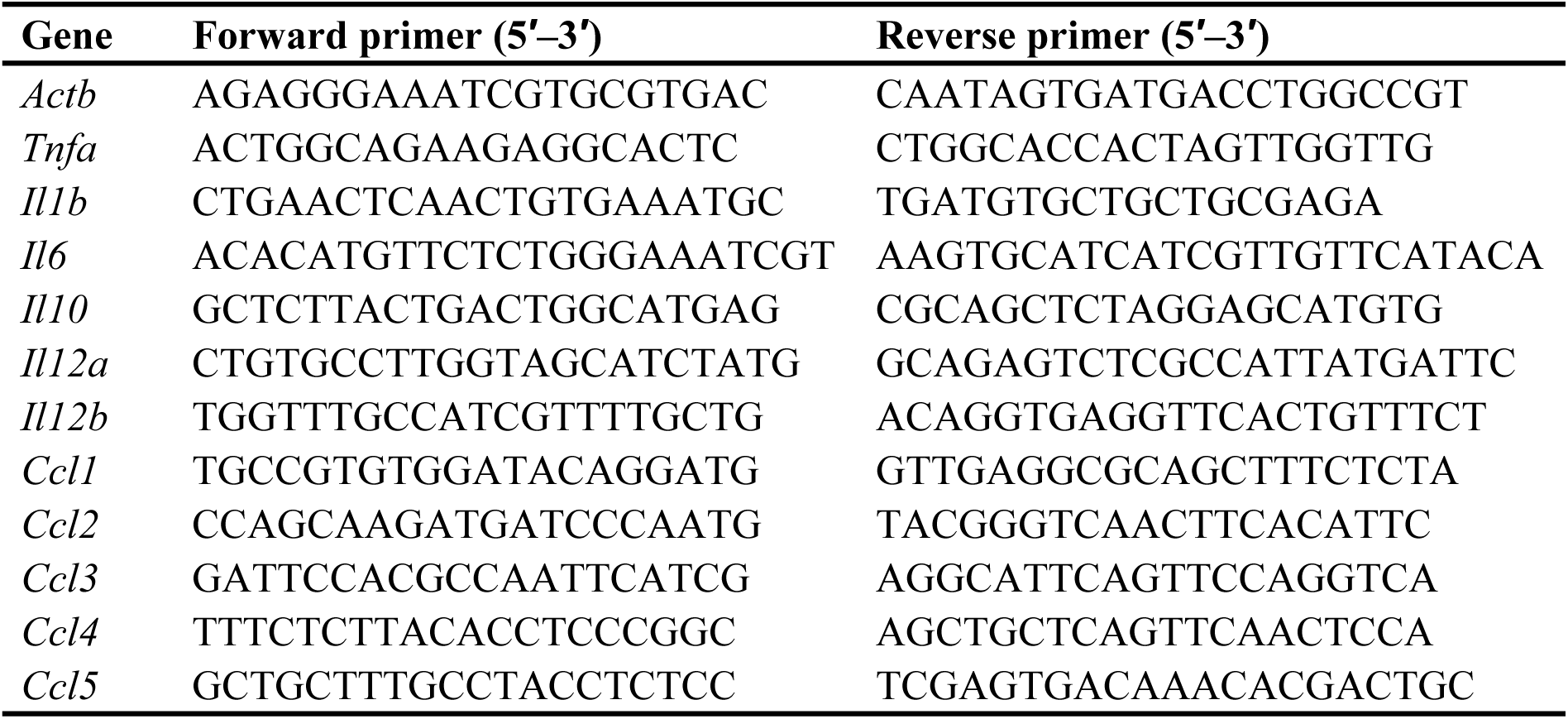
Sequences of primers

### Cytokine and chemokine assays

The protein levels of murine IL-10, CCL2, CCL3, CCL4, and CCL5 from whole liver and serum were measured using a cytometric bead assay flex set (BD Pharmingen) according to the manufacturer’s instructions. The levels of mouse IL-6, IL-12p40, and TNF-α were determined using a cytokine-specific enzyme-linked immunosorbent assay kit (R&D Systems). Protocols were used according to the manufacturer’s instructions.

### Analyses of serum ALT and hepatic hydroxyproline

For analysis of serum ALT, blood was sampled and centrifuged for collecting sera. For analysis of hepatic hydroxyproline, liver tissue was homogenized with an equal volume of PBS that contained a protease inhibitor cocktail (Sigma-Aldrich, St. Louis, Mo.) and centrifuged for collecting supernatants. Serum ALT and hepatic hydroxyproline levels were measured using commercially available kits (Jiancheng, Nanjing, China).

### Liver histology

A portion of the liver tissue was fixed in 4% paraformaldehyde, embedded in paraffin, cut into 5-μm sections, and stained with hematoxylin-eosin or picrosirius red using standard protocols.

### Statistical analysis

All data are expressed as mean ± SD and were analyzed using GraphPad Prism 6.01 software. Two-tailed, unpaired Student’s t tests were used to compare variables between two groups. *P* < 0.05 was considered statistically significant.

## Acknowledgements

We thank Professor Zhigang Tian for providing the *Il10*^−/−^ mice.

## Supporting information

**S1 Fig Quantitation of hepatic egg deposition in *Schistosoma japonicum*–infected WT mice and μMT mice.** Data represent mean ± SD; n = 5–7 samples per time point from two independent experiments. Two-tailed, unpaired Student’s *t* test.

**S2 Figure There is no difference in the numbers of circulating Ly6C^hi^ monocytes in peripheral blood of WT mice and μMT mice.**

(**A**) Gating strategy for detection of peripheral Ly6C^hi^ monocytes. (**B**) Representative flow cytometry plots of Ly6C^hi^ monocytes in peripheral blood of WT mice and μMT mice. (**C**) graphical summary showing percentage of peripheral Ly6C^hi^ monocytes out of total monocytes (left panel) and number of peripheral Ly6C^hi^ monocytes (right panel) in WT mice and μMT mice without infection (Ctrl) and 6 weeks after *Schistosoma japonicum* infection. Data represent mean ± SD; n = 3–5 per group from one experiment. \**p* < 0.05, two-tailed, unpaired Student’s *t* test.

**S3 Fig Serum chemokine levels in μMT mice are lower than those in MT mice.** Serum protein levels of CCL2, CCL3, CCL4 and CCL5 in WT mice and μMT mice were examined 6 weeks after *Schistosoma japonicum* infection. Data represent mean ± SD; n = 5–7 per group from two independent experiments. \**p* < 0.05, two-tailed, unpaired Student’s *t* test.

**S4 Fig Gating strategies for liver and PC B cell subsets.** (**A**) Representative flow cytometry plots show the gating strategy to identify hepatic B1a cells (CD3^−^CD19^+^CD5^+^CD23^−^IgM^hi^IgD^lo^), B1b cells (CD3^−^CD19^+^CD5^−^CD23^−^IgM^hi^IgD^lo^), and B2 cells (CD3^−^CD19^+^CD5^−^CD23^+^IgM^lo^IgD^hi^). (**B**) PC B1a cells were identified as CD3^−^CD19^+^CD5^+^CD11b^+^. PC B1b cells were identified as CD3^−^CD19^+^CD5^−^CD11b^+^. PC B2 cells were identified as CD3^−^CD19^+^CD5^−^CD11b^−^.

**S5 Fig The purity of hepatic B cells after sorting.** Representative flow cytometry plots showing the purity of sorted hepatic B cells derived from WT mice.

